# A Non-Alchemical Absolute Binding Free Energy Framework for Small Molecule Drugs

**DOI:** 10.64898/2026.02.18.706686

**Authors:** Yu Shi, Jianing Li

**Affiliations:** Borch Department of Medicinal Chemistry and Molecular Pharmacology, Purdue University, West Lafayette, IN 47907

**Keywords:** binding free energy, small molecule drug, molecular dynamics, solvent

## Abstract

A fully physical, non-alchemical framework is presented for absolute binding free energy (ABFE) calculations for protein–ligand complexes. Incorporation of a regularization potential eliminates unphysical artifacts and provides several key advantages: no endpoint catastrophes, fast convergence, robust performance for charged and neutral ligands, and rapid upfront verification. Validated on 30 diverse systems, the method improves predictive accuracy by 15.6% and numerical stability by 17.1% over leading alchemical approaches. This non-alchemical ABFE framework can be a potentially accurate and robust tool, carrying a potential for future computer aided small-molecule drug design.

## Introduction

Absolute binding free energy (ABFE) is a central quantity for quantifying drug–target molecular affinity in modern computational drug discovery. Among existing approaches, free energy perturbation (FEP) within an alchemical framework has become the dominant methodology for ABFE calculations in both academic and industrial settings. The most widely used protocol is the standard double-decoupling method (DDM), which constructs a thermodynamic cycle connecting two physical end states: one in which the ligand is fully solvated in bulk water, and another in which the ligand is fully solvated in the protein environment, regardless of whether it resides in a binding pocket or in the vicinity of the protein surface.

Within this alchemical framework, the absolute binding free energy is defined as the free energy difference between these two end states. Practically, this difference is obtained by decomposing the process into a sequence of alchemical transformations. First, the ligand is gradually decoupled from the bulk solvent by turning off its interactions with water molecules. This decoupling is typically performed in stages: electrostatic interactions are scaled down first, followed by the van *der* Waals interactions to avoid numerical instabilities. The resulting non-interacting (dummy) ligand is then transferred from the bulk solvent into the protein environment at no free energy cost. Subsequently, the ligand–protein interactions are restored in reverse order, with van *der* Waals interactions turned on first, followed by electrostatics. Throughout these steps, ligand–environment interactions are modulated using a series of λ windows, where the coupling parameter λ smoothly varies from 0 (fully decoupled) to 1 (fully interacting).

In this work, we introduce an ABFE framework distinct from the alchemical framework, providing accurate estimation for small molecule drugs. Although this alchemical scheme has been broadly adopted and validated, it suffers from a fundamental limitation — the presence of unphysical intermediate states. These partially interacting or non - interacting ligand configurations have no direct physical counterpart, which makes them difficult to reconcile with high-level quantum chemistry methods and quantum-informed or quantum-powered AI potential models. As a result, integrating alchemical free energy approaches with next-generation quantum mechanical descriptions of molecular interactions remains highly challenging. This intrinsic incompatibility constitutes a major bottleneck for further methodological advances and is the primary motivation for the development of a non-alchemical framework, in which free energy differences are evaluated exclusively through physically meaningful states and interactions.

### Theory, Methods, and Models

This all-physics ABFE innovation is illustrated by a detailed thermodynamic cycle that break down the ABFE calculation into nine phases (Figure 1). Specifically, rather than scaling solute-solvent interactions through non-physical intermediate states, this framework gradually introduces or decouples a regularization potential around the physical ligand (solid diamond) and the dummy ligand (empty diamond), as in Figure 1.

**Figure 1.**
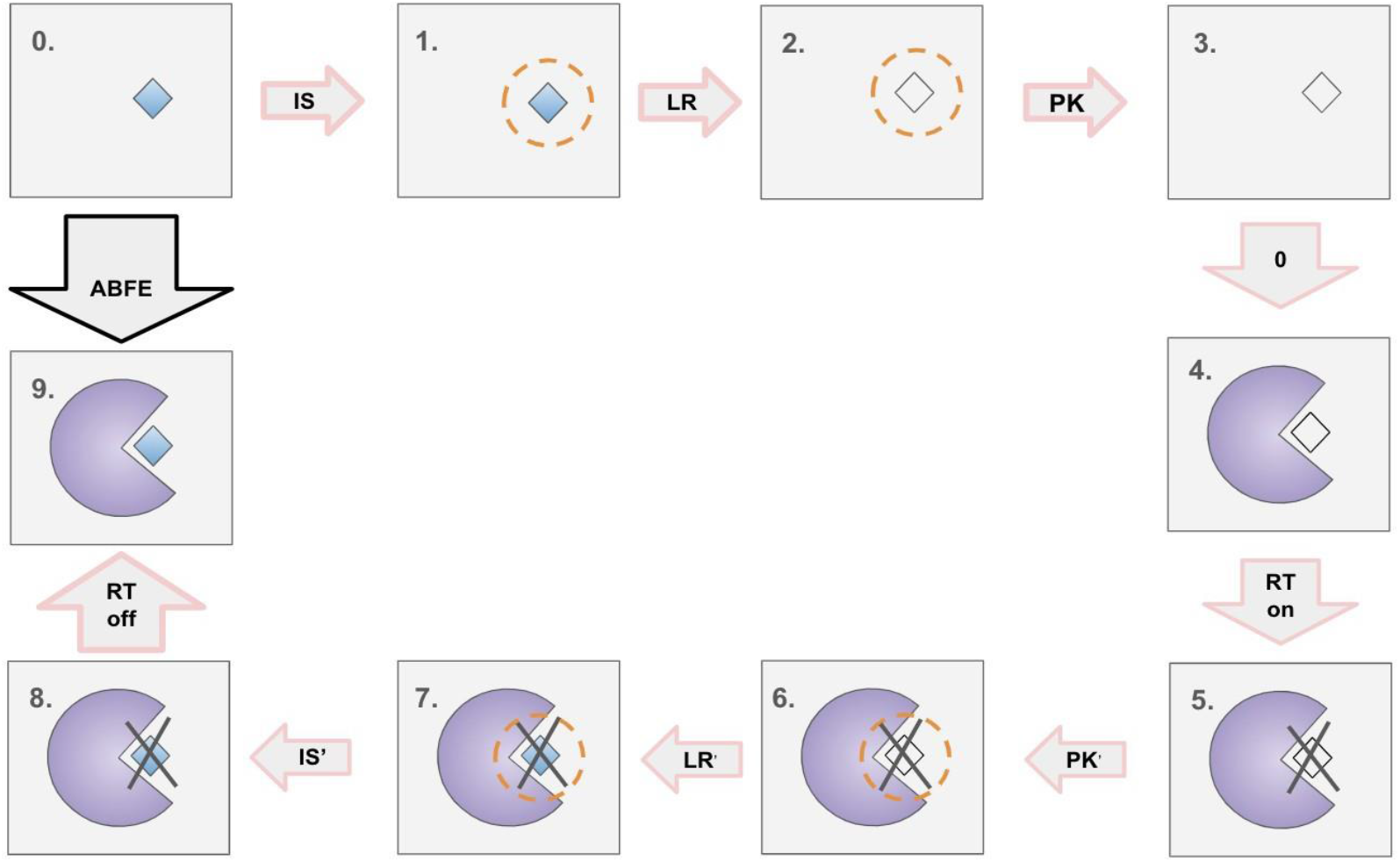
The Thermodynamic Cycle of the All-physics framework. The absolute binding free energy is the ligand free energy difference between the phase 0 and phase 1. The solid diamond represents the ligand fully interaction with solvents; the empty diamond repents the dummy ligand, which interactions with solvents are fully turned off. The purple circle presents the protein, and the grey area in each plot is for the bulk waters. The dash circle represents the regularization potential around the ligand solute. The cross indicates that there is restraint between the ligand and protein.

The physical ligand represents the state where ligand-solvent interactions are fully active, whereas the dummy ligand represents the state where these interactions are completely deactivated. The absolute binding free energy of the ligand to the protein is determined by quantifying the solvation free energy difference between Phase 0 (the physical ligand in a bulk water environment) and Phase 9 (the physical ligand bound to the protein environment). The thermodynamic cycle proceeds in two primary pathways: the top pathway describes the desolvation of the ligand from bulk water, while the bottom pathway describes the solvation of the ligand into the protein binding site. As defined in Eq. 1, the total solvation free energy is partitioned into three distinct physical contributions, including the inner-shell (IS) 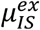, long-range (LR) 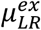, and packing (PK) term 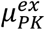.

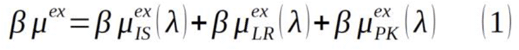

Further theoretical derivations and the rigorous physical basis for this partitioning scheme are detailed in previous research by Shi and Beck.^1^ Transitioning from Phase 0 to Phase 1, the regularization potential is introduced incrementally as the scaling factor γ increases from 0 to 1. In Phase 1, the fully activated regularization potential represented by the dashed circle surrounding the physical ligand, defines the coupled state.This regularization potential is based on the Weeks-Chandler-Andersen (WCA) potential as defined in Eq. 2, where λ represents the characteristic size of the potential. During this transition, the regularization is modulated via the scaling function f _λ_(γ), as specified in Eq. 3. The resulting free energy difference between Phase 0 and Phase 1 is thus defined as the IS contribution.

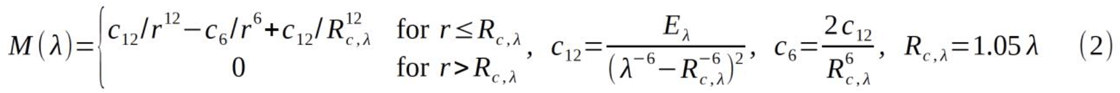

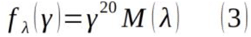

Following the initial coupling, the transition from Phase 1 to Phase 2 involves deactivating all ligand-solvent interactions except for the fully activated regularization potential. The resulting free energy change is defined as the LR contribution, and Phase 2 represents the uncoupled state. Subsequently, moving from Phase 2 to Phase 3, the regularization potential is annihilated stepwise as the scaling factor γ decreases from 1 to 0. This free energy difference is defined as the PK contribution. Notably, as shown in Phase 4, no free energy change occurs when the dummy ligand is transferred from the bulk water environment into the protein binding site. To maintain the ligand’s orientation and proximity within the binding site, Boresch restraints are applied.^2^ These restraints involve six geometric terms defined by three selected ligand atoms (A, B, C) and three protein atoms (a, b, c):

1. One harmonic distance potential between atoms A and a;
2. Two harmonic angle potentials for (a, b, A) and (a, A, B);
3. Three harmonic dihedral angle potentials for (c, b, a, A), (b, a, A, B), and (a, A, B, C).

The cycle continues from Phase 5 to Phase 6, where the regularization potential is reintroduced stepwise. In Phase 7, ligand-protein interactions are fully activated. From Phase 7 to Phase 8, the regularization potential is again annihilated. Finally, the Boresch restraints are removed through a sequence of scaling windows.

The primary innovation of this framework lies in its treatment of regularization potentials rather than direct scaling of complex ligand-solvent interactions. By avoiding non-physical artifacts (such as the ‘endpoint singularity’ problem in traditional FEP), this method preserves the physical integrity of each intermediate state. This characteristic significantly streamlines integration with high-level quantum chemistry (e.g., Density Functional Theory/DFT) or Quantum-AI potentials, as demonstrated in sodium ion solvation^3^and machine learning-based ion-pair solvation.^4^

To validate the predictive accuracy of this innovative framework, extensive molecular dynamics (MD) simulations were conducted on 30 ligand-protein complexes and 30 single-ligand systems. Initial structures for all systems were sourced from established peer-reviewed publications and public structural databases. The OpenMM software package^5^ was employed as the primary engine for all MD simulations. The standard computational protocol for each system follows five steps. The first step is to prepare the molecular models and input parameters. For proof of principle, this work adopted all the initial conformations from prior publications^6-8^ and the conformations were converting to the OpenMM format using Gromacs. As a repulsive potential between the heavy atoms on the ligand molecule and heavy atoms all of the solvent molecules, the regularization potential is added in the second step, with the scaling potential fλ(γ) as given in Eq. 3 as well as the Boresch restraints. The parameters for the Boresch restraints are from Ref. 6-7. With minimization in the third step, the fourth step is a 1-ns equilibration phase performed in the NVT ensemble using the Langevin Middle Integrator set at 300 K with a collision frequency of 1.0 ps^-1^ and a 2-fs step size for 500000 steps. Finally, a 5 ns production simulation was conducted in the NPT ensemble. A constant pressure was maintained using the Monte Carlo Barostat at 1 bar and 300 K.

To ensure statistical convergence and robust sampling, three independent replicas were generated for each system, with the results reported as the ensemble average. Each replica is independently equilibrated and uses the same Boresch restraint parameters. As illustrated in the Fig. 2 for the case-study Bromosporine Bound to CECR2, (PDB:3NXB), we employed Thermodynamic Integration (TI), as defined in Eq. 4 and Eq. 5, to compute the packing (PK) and inner-shell (IS) contributions across multiple γ windows. A step size of Δγ = 0.02 was utilized for all transitions. For the packing contribution, γ was varied from 0.20 to 1.00. For the inner-shell contribution, γ spanned from 0.40 to 1.00. These lower bounds were selected because the integrand effectively vanishes as γ approaches 0.20 and 0.40, respectively.

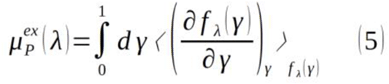

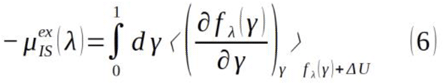

**Figure 2.**
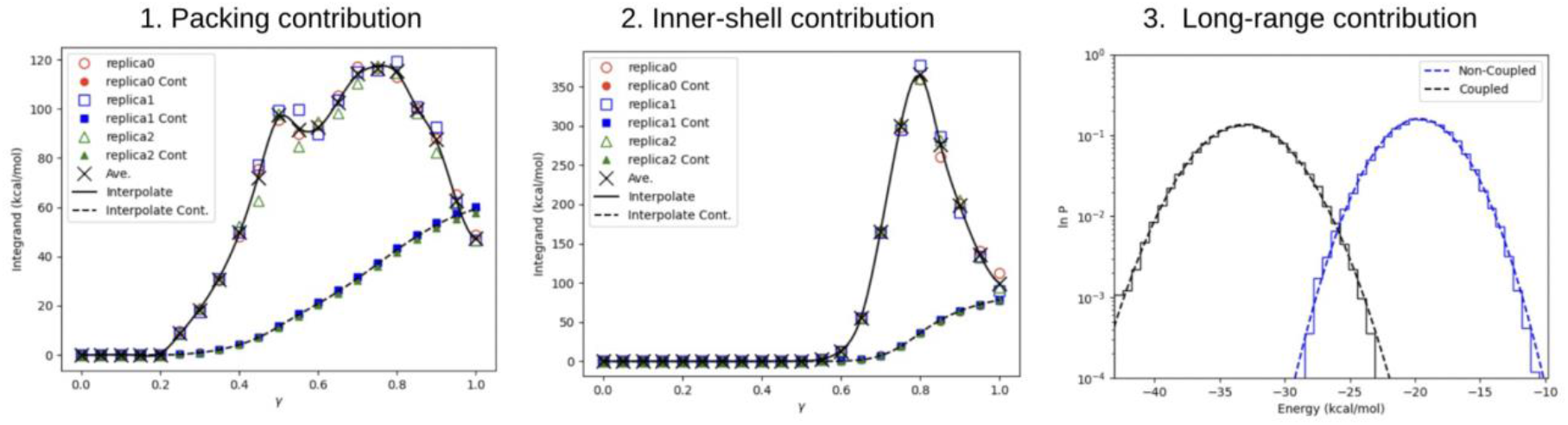
Computational analysis of Packing (PK), Inner-Shell (IS), and Long-Range (LR) contributions. For the packing and inner-shell terms, results from three independent replicas are plotted to demonstrate statistical consistency. In the long-range contribution plot, the ligand-solvent interaction energies for the coupled and uncoupled states are shown for one replica only, ensuring visual clarity. The interpolate results for the integrand and integration are shown as solid and dash curves. The Ave. with the cross symbol is the average value over three replicas.

In Eq. 6, the ΔU is the ligand-solvent interaction energy. A significant advantage of this framework is that when the scaling factor γ is zero, there is literally no contribution to the free energy integration. This feature allows the method to inherently bypass the “endpoint catastrophe”, a notorious numerical instability inherent in traditional alchemical frameworks when particles are created or annihilated. As illustrated in Fig. 2, our approach ensures a substantial phase-space overlap between the ligand-solvent interaction energies sampled from the coupled and uncoupled states. This overlap facilitates a robust and reliable estimation of the LR contribution by applying the near-Gaussian approximation as defined in Eq. 7.

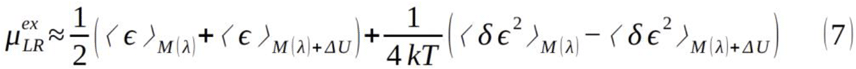

where ⟨*ϵ*⟩_*M*(*λ*)_ and ⟨*ϵ*⟩_*M*(*λ*)+Δ*U*_are the average interaction energy in the uncoupled and coupled states, ⟨*δϵ*^2^⟩_*M*(*λ*)_ and ⟨*δϵ*^2^⟩_*M*(*λ*)+Δ*U*_ are the standard deviations of interaction energy in the uncoupled and coupled, k is the Boltzmann constant, T is the temperature 300 K.

For the removal the Boresch restraint potentials, 10 non-uniformly distributed γ windows were used (0.000, 0.010, 0.025, 0.050, 0.075, 0.100, 0.250, 0.500, 0.750, 1.000). A total 38 windows were therefore employed.

As shown in the Ref. 1, the solute solvation free energy should be independent of the regularization potential shape or sizes we use. This is supported by data shown in Table 1, where two simulations were run with two different regularization potential sizes, one is 4.5 Å and the other one is 5.0 Å. Each individual contribution is quite different between the simulations, however as the sum of them (indicated by the last column “Sol.”), the ligand solvation free energies are highly consistent. This provides us with an important advantage to run a quick verification upfront before committing to any large-scale simulations. Further analysis demonstrated (Figure 3) that each individual free energy contribution exhibits remarkably rapid convergence. While our standard protocol utilizes a 5 ns production simulation, a reliable and statistically significant prediction is typically achieved within the first nanosecond. This accelerated convergence represents a transformative shift in computational efficiency. The ability to achieve convergence in a fraction of the standard simulation time may reduce the wall-clock time and operational costs associated in drug discovery.

**Table 1.** the independence of regularization potential (R_Potential) Size.

| R_Potential Size ($\text{\AA}$) | PK/std. (kcal/mol) | IS/std. (kcal/mol) | RL/std. (kcal/mol) | Sol./std. (kcal/mol) |
| --- | --- | --- | --- | --- |
| 4.5 | 62.00/0.15 | -76.65/1.70 | -173.84/0.41 | -188.49/1.76 |
| 5.0 | 74.36/0.98 | -99.27/0.65 | -164.49/0.39 | -189.40/1.23 |

**Figure 3.**
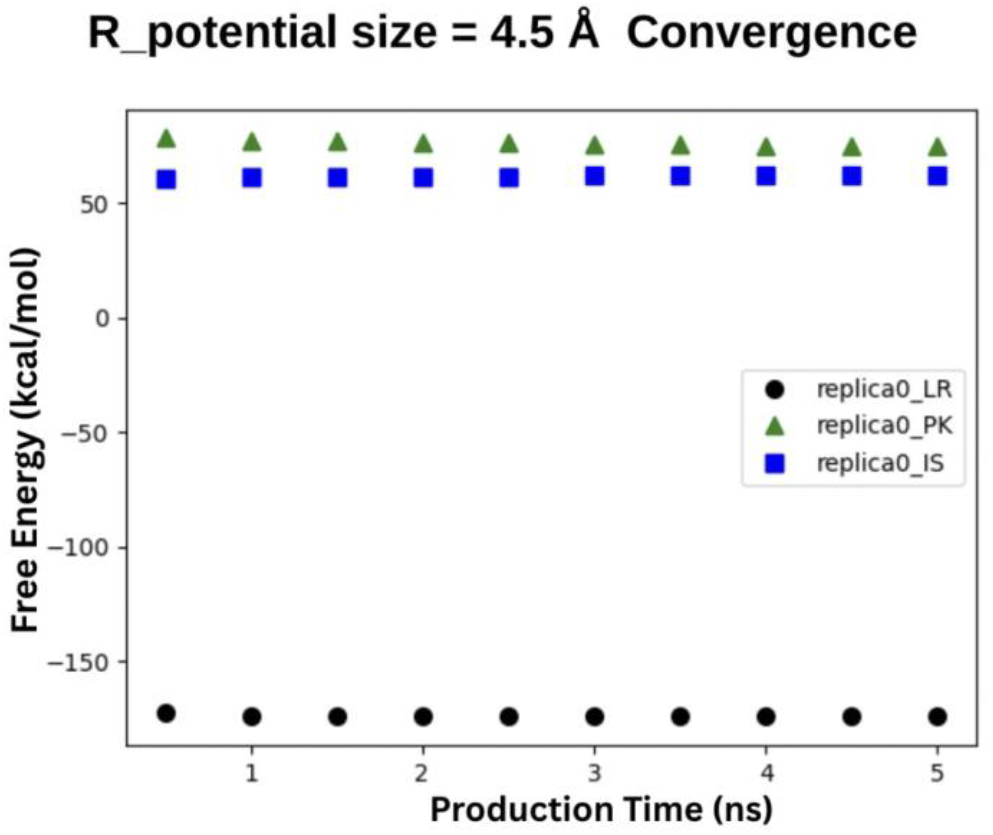
The convergence of each individual contribution with production simulation time. The triangle is for the packing contribution, the circle point is the long-ranged contribution, and the square is for the inner-shell contributions. The simulation is with the regularization potential size 4.5 Å for the Case Study Bromosporine Bound to CECR2. For clarity, only the result from one replica is presented.

## Results and Discussions

In total, 30 protein-small molecule complex systems with binding energy from -4 to -12 kcal/mol (Table 3) were tested with this ABFE framework, yielding 16% improvement in the accuracy and 17% in the robustness (Table 2 and 3), suggesting an improvement over the traditional alchemical framework. In Figure 4, the shaded gray areas indicate where the 1 and 2 kcal/mol error boundaries lie. The Pearson’s rp = 0.90, while Spearman’s rs = 0.93. In the square brackets are the 95% confidence intervals (CI) of the statistics based on percentile bootstrap. The solid line represents the linear regression, demonstrating a predictive slope of 0.94 (Ideal 1.0 as shown by the dashed line). All these data indicate a good agreement between our calculations and reference experimental values.

**Table 2.** Comparison of MUE and RMSE between the calculations by Non -Alchemical and Alchemical framework. In the square brackets are the 95% confidence intervals (CI) of the statistics based on percentile bootstrap.

| Metrics | Non-Alchemical (kcal/mol) | Alchemical (kcal/mol) | Improvement |
| --- | --- | --- | --- |
| MUE (Accuracy) | 0.65<br>[0.48,0.84] | 0.77<br>[0.56,1.00] | 15.6% |
| RMSE (Robustness) | 0.82<br>[0.61,1.04] | 0.99<br>[0.71,1.28] | 17.1% |

**Table 3.** A summary of all the 30 complexes reported in this work. For each complex, the experimental data, alchemical data were compared with the ABFE (non -alchemical) results.

| Complex (protein/Ligand) | Experiment data (kcal/mol) | Non-Alchemical with OpenMM (kcal/mol) | Alchemical with Gromacs (kcal/mol) |
| --- | --- | --- | --- |
| Philip Biggin JACS 2022 (Others in Figure 4) Ref. 6 |  |  |  |
| PWWP1 / Ligand-8 | -5.2 | -4.3 | -4.5 |
| PWWP1 / Ligand-9 | -6.4 | -7.0 | -6.8 |
| PWWP1 / Ligand-10 | -6.5 | -7.3 | -6.1 |
| PWWP1 / Ligand-11 | -6.6 | -7.1 | -6.6 |
| HSP90 / Ligand-3 | -4.2 | -5.2 | -5.1 |
| MCL / Ligand-17 | -5.7 | -5.9 | -5.5 |
| Cyclophilin / Ligand2 | -9.0 | -9.4 | -8.1 |
| NPGN/T4L / L99A | -5.5 | -6.2 | -4.8 |
| Philip Biggin JACS 2017 Ref. 7 |  |  |  |
| BRD2_2V / RVXOH | -8.8 | -9.0 | -7.9 |
| BRD3_1H / RVXOH | -8.0 | -7.9 | -6.7 |
| BRD3_2V / RVXOH | -8.8 | -8.5 | -7.4 |
| BRD4_1H / RVXOH | -9.0 | -10.3 | -9.9 |
| BRD4_2V / RVXOH | -8.8 | -9.0 | -8.5 |
| BRDT_1H / RVXOH | -7.7 | -8.6 | -7.7 |
| BRD2_1V / RVX208 | -6.9 | -6.9 | -7.2 |
| BRD2_2V / RVX208 | -8.5 | -8.0 | -7.5 |
| BRD3_1V / RVX208 | -7.1 | -7.9 | -5.5 |
| BRD3_2V / RVX208 | -8.8 | -10.0 | -8.7 |
| BRD4_1V / RVX208 | -7.8 | -7.5 | -6.8 |
| BRD4_2V / RVX208 | -9.1 | -8.5 | -9.7 |
| BRDT_1V / RVX208 | -7.0 | -7.6 | -6.5 |
| CECR2 / Bromosporine | -10.7 | -12.1 | -11.5 |
| BRD2_2 / Bromosporine | -9.6 | -11.8 | -9.7 |
| BRD3_1 / Bromosporine | -9.1 | -9.1 | -7.2 |
| BRDT_1 / Bromosporine | -9.8 | -9.1 | -9.3 |
| BRPF1B / Bromosporine | -11.5 | -9.8 | -8.6 |
| TIF1 / Bromosporine | -6.7 | -7.0 | -5.4 |
| TAF1_2 / Bromosporine | -10.2 | -10.2 | -10.7 |
| PB1_5 / Bromosporine | -6.4 | -6.9 | -7.0 |
| SMAR. / Bromosporine | -6.2 | -6.9 | -7.0 |

**Figure 4.**
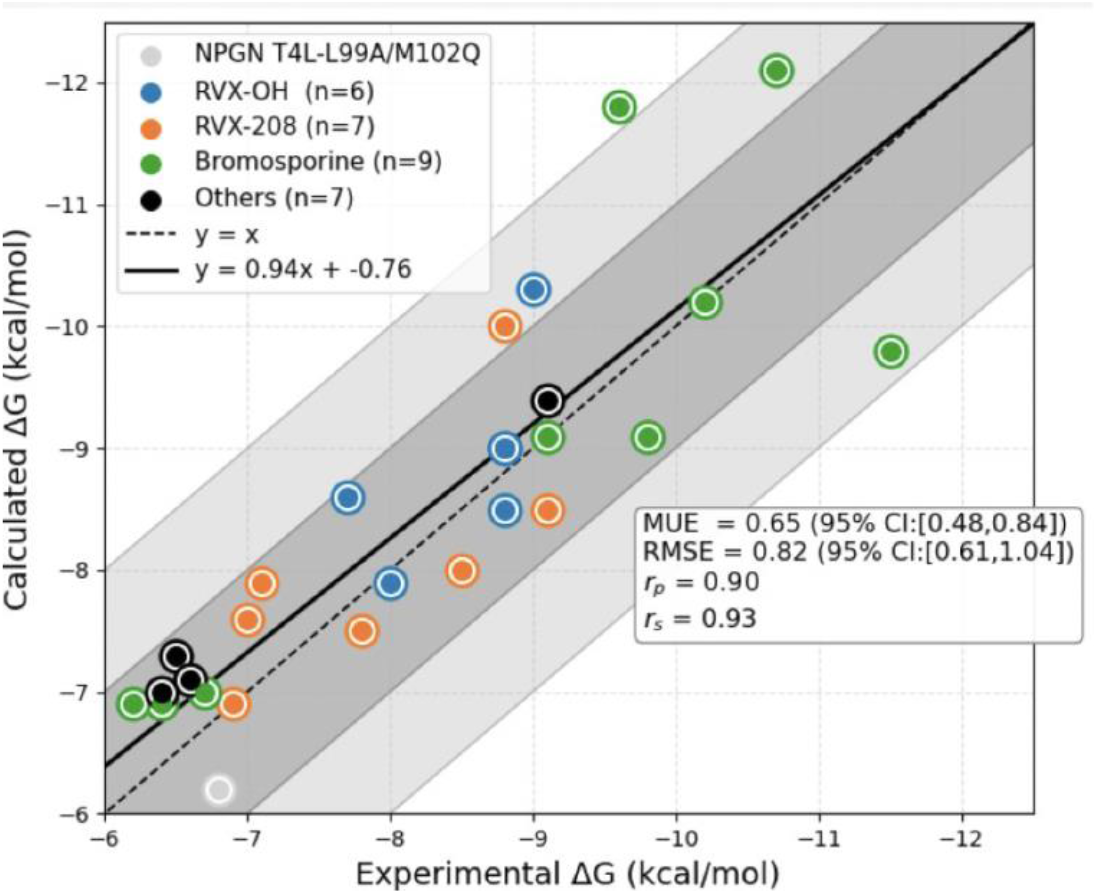
Scatter plot of calculated vs experimental affinities for 30 protein -ligand complexes. The grey dot (NPGN-T4-L99A/M102Q) is n-phenylglycinonitrile combined with engineered T4 lysozyme cavity (Online tutorial ABFE Gromacs 2016, Ref. 8). Complexes with ligand of RVX-OH, RVX-208 and Bromosporine are Biggin’s JACS 2017 paper, Ref. 7. Others are from Ref. 8. The shaded gray areas indicate where the 1 and 2 kcal/mol error boundaries lie. The Pearson’s rp = 0.90, while Spearman’s rs = 0.93. In the square brackets are the 95% confidence intervals (CI) of the statistics based on percentile bootstrap. The solid line represents the linear regression, demonstrating a predictive slope of 0.94 (Ideal 1.0 as shown by the dashed line)

Notably, these results represent the inaugural implementation of this framework. The current performance was achieved using classical force fields for proof of principle. There remains significant room for further optimization in the regularization potential’s functional form and its coupling parameters. As we transition to more advanced AI-driven potentials and refined sampling strategies, we anticipate that the predictive power and efficiency of this framework will see even more substantial gains in the near future.

## Conclusion

This work describes a non-alchemical ABFE framework as a potentially accurate and robust tool, carrying a potential for future computer aided small-molecule drug design. Current tests with a data set of 30 complexes provides proof of principle to support the future larger-scale applications.

## Acknowledgements

We thank Dr. Tom Beck (Oakridge National Lab) and Dr. Wei Jiang (Argonne National Lab) for helpful discussions. This work was supported by the AnalytiXIN fellowship to J. L. and the computational resources provided by Purdue’s RCAC and Argonne’s ALCF.

## References

1. Yu Shi and Thomas Beck, The Journal of Chemical Physics 139, 044504 (2013).

2. Stefan Boresch, Journal of Chemical Information and Modeling 64, 3605–3609 (2024).

3. Yu Shi and Thomas Beck, Proceedings of the National Academy of Sciences 117 (48) 30151 (2020)

4. Yu Shi, Stephen T. Lam and Thomas L. Beck, Chemical Science 13 8265, (2022)

5. Eastman, R. Galvelis, R. P. Peláez, C. R. A. Abreu, S. E. Farr, E. Gallicchio, A. Gorenko, M. M. Henry, F. Hu, J. Huang, A. Krämer, J. Michel, J. A. Mitchell, V. S. Pande, J. PGLM Rodrigues, J. Rodriguez-Guerra, A. C. Simmonett, S. Singh, J. Swails, P. Turner, Y. Wang, I. Zhang, J. D. Chodera, G. De Fabritiis, and T. E. Markland. “OpenMM 8: Molecular Dynamics Simulation with Machine Learning Potentials.” Journal of Physical Chemistry B 128(1), pp. 109–116 (2023).

6. Irfan Alibay, Aniket Magarkar, Daniel Seeliger & Philip Charles Biggin, Communications Chemistry, volume 5, Article number: 105 (2022)

7. Matteo Aldeghi, Alexander Heifetz, Michael J Bodkin, Stefan Knapp, Philip C Biggin, Journal of the American Chemical Society, 139(2):946–957 (2017)

8. Online tutorial, Absolute Binding Free Energy - Gromacs 2016, https://www.alchemistry.org/wiki/Absolute_Binding_Free_Energy_-_Gromacs_2016

